# The Ribosomal Operon Database (ROD): A full-length rDNA operon database derived from genome assemblies

**DOI:** 10.1101/2024.04.19.590225

**Authors:** Anders K. Krabberød, Embla Stokke, Ella Thoen, Inger Skrede, Håvard Kauserud

## Abstract

Current rDNA reference sequence databases are tailored towards shorter DNA markers, such as parts of the 16/18S marker or the ITS region. However, due to advances in long-read DNA sequencing technologies, longer stretches of the rDNA operon are increasingly used in environmental sequencing studies to increase the phylogenetic resolution. There is, therefore, a growing need for longer rDNA reference sequences. Here, we present the Ribosomal Operon Database (ROD), which includes eukaryotic full-length rDNA operons fished from publicly available genome assemblies. Full-length operons were detected in 34.1% of the 34,701 examined eukaryotic genome assemblies from NCBI. In most cases (53.1%), more than one operon variant was detected, which can be due to intragenomic operon copy variability, allelic variation in non-haploid genomes, or technical errors from the sequencing and assembly process. The highest copy number found was 5,947 in *Zea mays*. In total, 453,697 unique operons were detected, with 69,480 operon variant clusters remaining after intragenomic clustering at 99% sequence identity. The operon length varied extensively across eukaryotes, ranging from 4,136 to 16,463 bp, which will lead to considerable PCR bias during amplification of the entire operon. Clustering the full-length operons revealed that the different parts (i.e., 18S, 28S, the hypervariable region V4 of 18S, and ITS) provide divergent taxonomic resolution, with 18S and the V4 region being the most conserved. The Ribosomal Operon Database (ROD) will be updated regularly to provide an increasing number of full-length rDNA operons to the scientific community.

## Introduction

DNA metabarcoding, high-throughput sequencing of short DNA regions amplified from environmental DNA followed by taxonomic annotation using known reference sequences, is a powerful approach to explore microbial communities (Taberlet et al., 2018). Different parts of the ribosomal rDNA operon are the predominant markers used in DNA metabarcoding studies, such as the V4 region commonly used for microeukaryotes (Hadziavdic et al., 2014; Krabberød et al., 2022; Logares et al., 2012) or the ITS regions commonly used for fungi (Schoch et al., 2012). In the last decade and a half, relatively short DNA markers within the 200-400 bp range have typically been used, mainly due to inherent length limitations of relevant sequencing technologies. For taxonomic annotation of these markers, designated sequence databases such as PR2 (Guillou et al., 2013), RDP (Maidak et al., 2001), SILVA (Quast et al., 2012), and UNITE (Nilsson et al., 2019) have been constructed, including millions of relatively short rDNA reference sequences.

In contrast to short-read DNA metabarcoding, long-read sequencing technologies make it possible to sequence longer stretches of the rDNA region (Jamy et al., 2020), even the entire rDNA operon (Wurzbacher et al., 2019), with high precision (Callahan et al., 2019). This calls for new reference sequences covering the whole rDNA operon. Comprehensive full-length operon databases have been developed for prokaryotes (Kerkhof et al., 2022; Seol et al., 2022) but not so far for eukaryotic organisms. One way to construct full-length operon reference sequences is to extract this information from genome assemblies; with increasing speed and precision, genome assemblies are becoming publicly available, driven by large-scale initiatives such as the Earth Biogenome Project (Lewin et al., 2022), ERGA (Formenti et al., 2022), Mycocosm (Grigoriev et al., 2014) and Darwin Tree of Life (Darwin Tree of Life Project Consortium, 2022). However, finding and extracting complete, full-length operons from genome assemblies can be difficult since technical errors may occur during sequencing and bioinformatics processing, rendering the ribosomal operon incomplete. Moreover, since the rDNA region is a multi-copy marker, intragenomic variants can exist (Lindner et al., 2013; Simon & Weiss, 2008) due to incomplete homogenization of the repeats (Elder & Turner, 1995; Ganley & Kobayashi, 2007).

Using longer stretches of the rDNA operon as a DNA metabarcoding marker can also pose extra challenges during PCR, partly due to its length, making it harder for the polymerase enzyme to read and copy the entire region. Moreover, various PCR biases, such as operon length differences among the target taxa, can skew the results. Although not thoroughly studied across the Eukaryotic Tree of Life, a considerable length variation occurs between lineages (Hall et al., 2022). One of the main reasons for this variation is introns, which have been shown to be prevalent in both 18S and 28S (Bhattacharya et al., 2000; Cannone et al., 2002; Hedberg & Johansen, 2013; Karpov et al., 2017).

Different parts of the rDNA operon mutate and evolve at different rates (Clark, 1987; Hall et al., 2022; Hillis & Dixon, 1991). Compared to the relatively conserved 18S and 28S regions, the ITS region evolves rapidly and can provide highly resolved taxonomic information down to the species level (Kauserud, 2023; Schoch et al., 2012). Different parts of the 18S and 28S genes also evolve at varying rates based on their position in the final rRNA molecule; in 18S, there are nine regions known to be hypervariable (named V1 to V9) (Hadziavdic et al., 2014; Nelles et al., 1984; Rajilić-Stojanović et al., 2009), while in 28S, there are twelve named D1-D12 (Chilton et al., 2003; Hassouna et al., 1984). These regions have different lengths and variability.

For instance, the V4 region in 18S shows significant sequence variability (Hadziavdic et al., 2014) and has been extensively used as a marker in DNA metabarcoding projects (Hadziavdic et al., 2014). D1, D2, and D3 of the 28S have similar properties and have been used to identify species where 18S has not provided the required taxonomic resolution (Douda et al., 2013; Qing et al., 2017; Rué et al., 2023; Sonnenberg et al., 2007). Knowing the level of variability and which taxonomic resolution the markers provide is crucial in DNA metabarcoding analyses. However, we have limited knowledge about the relative resolution of the different parts of the rDNA operon and how this may vary between taxonomic groups.

In this study, we introduce the full-length Ribosomal Operon Database (ROD) constructed from publicly available genome assemblies of eukaryotes. In addition to providing a high number of full-length operon reference sequences for the scientific community, we report on different features of the rDNA operons, including multiple operon variants in many genomes and length differences across eukaryotes. We also compare the level of molecular variation and how this affects the number of sequence clusters across different parts of the rDNA operon (18S, 28S, V4, ITS1, and ITS2), as well as the entire operon.

## Materials and methods

### Fishing reference operons

Initially, Barrnap (Seemann, n.d.) was used to identify the ribosomal RNA genes 18S, 5.8, and 28S from a selection of 947 genomes from Phycocosm (Grigoriev et al., 2021) and Mycocosm (Grigoriev et al., 2014). Following the initial identification, the output from Barrnap was analyzed to manually extract genomic regions where 18S, 5.8S, and 28S were arranged as a continuous operon on the same contig or scaffold, and a new Hidden Markov Model (HMM) was built to represent the complete operon. This model was constructed with hmmer3 (*HMMER 3*.*4 (Aug 2023)*, n.d.) from a MAFFT (Katoh & Standley, 2013) alignment of 940 full-length ribosomal operons, representing a wide array of eukaryotic lineages and covering the major branches of the eukaryotic tree of life.

All eukaryotic genomes available in NCBI as of December 2023, in total 34,701, were downloaded and screened with the new full-length operon model using hmmscan. Regions matching the ribosomal operon were found in 17,107 of the genomes. To maintain the focus on complete, full-length operons, Barrnap (Seemann, n.d.) was used to search for operons with incomplete 18S or 28S; in the cases where either of these genes were marked as “partial,” the operons were excluded from further analysis. After the filtration step, we were left with 453,697 operon sequences that we could say with reasonable confidence were full length.

### Determining operon variants

We used VSEARCH (Rognes et al., 2016) to cluster full-length operons within each genome assembly at a 99% similarity threshold. This clustering aimed to eliminate potential technical errors from sequencing and assembly, and to identify operon variants that diverge by more than 1% in sequence identity. Operons that diverge by more than 1% in sequence identity are likely to represent biologically meaningful variants rather than just technical noise. Clustering at 99% allows these divergent operon variants to be distinguished and analyzed separately.

### Eliminating contaminants

Several measures were taken to eliminate contaminants and incorrectly annotated genomes. This was done by comparing the centroid representatives from the clustered operons to the best hits in NCBI GenBank using blastn (Altschul et al., 1990). For each operon, 50 hits were retrieved from the NCBI nucleotide database and sorted by bitscore, e-value, and identity. The top hit for each query was then selected.

Taxonomic information was standardized using the R package *taxonomizr* (Sherrill-Mix, 2023) for both the retrieved genomes and the blast hit. The taxonomy identifier (TaxId) from the genomes in NCBI and the best hits for each centroid were used in the function *getTaxonomy()*, and the option for desired taxonomic levels was set to retrieve *superkingdom, kingdom, phylum, subphylum, class, order, family, genus*, and *species*. A comparative analysis of taxonomic assignments revealed possible discrepancies between the genome annotations and blast results, particularly in the higher taxonomic levels from *superkingdom* down to *order*. Operons with conflicting taxonomic assignments at these levels were excluded from the database to maintain its integrity. Additionally, operons with incomplete taxonomic information were quality-controlled by manually examining the top 50 blast hits, and a taxonomic assignment was selected to determine whether it was the majority among those matches. A total of 8614 centroids representing 34,015 ribosomal operons from 2,500 genomes were removed as contaminants. For 1,075 genomes, we uncovered one or more contaminant operons in addition to the operon corresponding to the expected taxonomy of the sequenced genome.

### Streamlining taxonomy

Finally, the taxonomic assignments for the remaining operons were standardized to 9 different levels, following the setup in PR2 (Guillou et al., 2013). The levels used in ROD are *domain, supergroup division, subdivision, class, order, family, genus*, and *species*. The higher levels *domain, supergroup, division*, and *subdivision* correspond roughly to *superkingdom, kingdom, phylum*, and *subphylum* in NCBI but are adjusted to recent suggestions for the organization of the major branches of the Tree of Life (Adl et al., 2019; Burki et al., 2019). In addition to the taxonomic levels, ROD contains the GenBank *assembly_id* and the *taxid* from NCBI for each genome. After checking for length, clustering, and removing contaminants, the database comprised 69,480 operon variants, representing 453,697 copies of the ribosomal operon from 11,935 genomes.

### Genetic distance

The operon variants were aligned separately for each genome assembly with more than three variants using MAFFT (Katoh & Standley, 2013). After aligning, the genetic distance was calculated for each alignment using the *dist*.*dna()* function (Paradis & Schliep, 2019) with the TN93 model, which assumes distinct rates for both kinds of transition (A ↔ G versus C ↔ T) and transversions (Tamura & Nei, 1993), and the base rates were estimated from the data. The distance calculation was weighted by the geometric mean of the size of the variants, i.e.

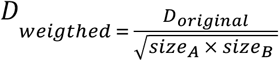

where *size*_*A*_ and *size*_*B*_ represent the copy number of two given operon variants and *D*_*original*_ the unweighted genetic distance between the two. The average genetic distance was then calculated for each subdivision on the Tree of Life (Table 1).

**Table 1.** Characteristics of rDNA operon in eukaryotes for each subdivision. Detailed statistics of ribosomal operons extracted from eukaryotic genomes in NCBI, categorized by Supergroup, Division, and Subdivision. Each entry provides data on the number of genomes analyzed for each subdivision, operon copy number statistics (mean and maximum), variant counts, average genetic distance, and operon length metrics (minimum, maximum, mean). The number of operon variants is defined as sequences at least 1% dissimilar, and the average genetic distance is calculated based on the variants and weighted by the number of copies per variant. *Genomes*: Number of genomes examined for the particular subdivision. *Avg. copy number*: Average number of ribosomal operon copies per genome in the subdivision. *Max Copies*: Maximum number of operon copies found in a single genome within the category. *Max Variants:* Maximum number of operon variants found among the genomes in the subdivision.*Avg. variants*: Average number of operon variants within the subdivision. *Min/Max/Avg Operon Length*: The minimum, maximum, and average length of operons within each subdivision, expressed in base pairs (bp). Cells containing ‘-’ indicate data not available or not applicable for specific categories.

| *Supergroup* | *Division* | *Subdivision* | *Genomes* | *Avg.*<br>*copy*<br>*number* | *Max copies* | *Max.*<br>*variants* | *Avg.*<br>*variants* | *Avg. genetic*<br>*distance* | *Min.*<br>*operon*<br>*length* | *Max.*<br>*operon*<br>*length* | *Avg.*<br>*operon*<br>*length* |
| --- | --- | --- | --- | --- | --- | --- | --- | --- | --- | --- | --- |
| Amoebozoa | Discosea | Discosea X | 10 | 59 | 11.3 | 36 | 6.9 | 0.006 | 4,814 | 7,027 | 5,765 |
| Amoebozoa | Evosea | Eumycetozoa | 7 | 2 | 1.1 | 1 | 1.0 | - | 5,260 | 6,837 | 6,362 |
| Amoebozoa | Evosea | Variosea | 2 | 1 | 1.0 | 1 | 1.0 | - | 4,891 | 5,778 | 5,335 |
| Amoebozoa | Tubulinea | Echinamoebida | 1 | 8 | 8.0 | 2 | 2.0 | 0.103 | 6,055 | 6,125 | 6,125 |
| Apusozoa | Apusozoa_XX | Apusozoa_XXX | 1 | 1 | 1.0 | 1 | 1.0 | - | 6,118 | 6,118 | 6,118 |
| Archaeplastida | Glaucophyta | Glaucophyta X | 1 | 5 | 5.0 | 1 | 1.0 | - | 5,381 | 5,381 | 5,381 |
| Archaeplastida | Rhodophyta | Rhodophyta | 29 | 21 | 5.7 | 10 | 3.2 | 0.006 | 4,694 | 6,634 | 6,073 |
| Archaeplastida | Viridiplantae | Streptophyta | 1,922 | 5,947 | 155.8 | 2,097 | 24.7 | 0.033 | 4,136 | 13,249 | 5,645 |
| Archaeplastida | Viridiplantae | Chlorophyta | 102 | 112 | 7.1 | 6 | 1.3 | 0.005 | 4,347 | 7,448 | 5,695 |
| CRuMS | Mantamonadidea | Mantamonadidea X | 1 | 3 | 3.0 | 1 | 1.0 | - | 6,527 | 6,527 | 6,527 |
| Excavata | Discoba | Euglenozoa | 84 | 36 | 4.8 | 11 | 1.5 | 0.001 | 4,684 | 7,448 | 6,627 |
| Excavata | Discoba | Heterolobosea | 7 | 6 | 1.9 | 1 | 1.0 | - | 5,605 | 6,105 | 5,766 |
| Excavata | Metamonada | Fornicata | 2 | 6 | 3.5 | 3 | 2.0 | 0.004 | 5,580 | 5,975 | 5,778 |
| Excavata | Metamonada | Preaxostyla | 1 | 17 | 17.0 | 2 | 2.0 | 0.003 | 4,757 | 6,047 | 4,757 |
| Haptista | Haptophyta | Haptophyta X | 3 | 20 | 9.7 | 1 | 1.0 | - | 4,915 | 6,020 | 5,535 |
| Opisthokonta | Filasterea | Filasterea X | 2 | 3 | 2.0 | 1 | 1.0 | - | 5,367 | 6,611 | 5,989 |
| Opisthokonta | Fungi | Ascomycota | 5,676 | 1,078 | 6.1 | 81 | 1.6 | 0.012 | 4,185 | 6,905 | 5,595 |
| Opisthokonta | Fungi | Basidiomycota | 772 | 463 | 8.9 | 26 | 1.6 | 0.003 | 4,272 | 6,691 | 5,632 |
| Opisthokonta | Fungi | Mucoromycota | 109 | 68 | 4.9 | 5 | 1.6 | 0.005 | 4,263 | 6,646 | 5,692 |
| Opisthokonta | Fungi | Zoopagomycota | 41 | 49 | 3.0 | 4 | 1.1 | 0.000 | 4,855 | 7,105 | 6,090 |
| Opisthokonta | Fungi | Chytridiomycota | 22 | 11 | 2.6 | 1 | 1.0 | - | 4,787 | 6,409 | 5,803 |
| Opisthokonta | Fungi | Blastocladiomycota | 3 | 8 | 5.3 | 2 | 1.3 | 0.001 | 4,486 | 6,058 | 6,013 |
| Opisthokonta | Fungi | Fungi_X | 1 | 1 | 1.0 | 1 | 1.0 | - | 4,706 | 4,706 | 4,706 |
| Opisthokonta | Fungi | Aphelidiomycota | 1 | 5 | 5.0 | 1 | 1.0 | - | 5,102 | 5,102 | 5,102 |
| Opisthokonta | Ichthyosporia | Ichthyosporia X | 2 | 4 | 2.5 | 4 | 2.5 | 0.006 | 5,407 | 5,486 | 5,446 |
| Opisthokonta | Metazoa | Arthropoda | 1,849 | 4,691 | 42.1 | 139 | 3.4 | 0.008 | 4,395 | 16,463 | 6,685 |
| Opisthokonta | Metazoa | Chordata | 442 | 1,007 | 38.3 | 665 | 5.6 | 0.015 | 4,162 | 13,305 | 5,370 |
| Opisthokonta | Metazoa | Nematoda | 132 | 448 | 23.3 | 15 | 2.1 | 0.007 | 4,415 | 10,248 | 5,972 |
| Opisthokonta | Metazoa | Mollusca | 108 | 505 | 24.9 | 96 | 3.7 | 0.027 | 4,640 | 11,390 | 6,384 |
| Opisthokonta | Metazoa | Cnidaria | 49 | 880 | 41.2 | 40 | 4.1 | 0.008 | 4,409 | 10,417 | 5,936 |
| Opisthokonta | Metazoa | Rotifera | 33 | 25 | 3.3 | 4 | 1.3 | 0.002 | 4,453 | 10,979 | 6,228 |
| Opisthokonta | Metazoa | Echinodermata | 26 | 710 | 58.9 | 45 | 3.4 | 0.014 | 4,772 | 10,310 | 5,865 |
| Opisthokonta | Metazoa | Annelida | 26 | 646 | 72.5 | 6 | 1.9 | 0.003 | 4,440 | 10,955 | 6,612 |
| Opisthokonta | Metazoa | Porifera | 18 | 258 | 47.0 | 8 | 2.6 | 0.009 | 4,624 | 7,588 | 5,960 |
| Opisthokonta | Metazoa | Platyhelminthes | 14 | 679 | 63.7 | 76 | 10.1 | 0.011 | 4,910 | 11,943 | 6,426 |
| Opisthokonta | Metazoa | Ctenophora | 4 | 4 | 2.0 | 2 | 1.3 | 0.016 | 4,558 | 6,236 | 5,275 |
| Opisthokonta | Metazoa | Tardigrada | 4 | 3 | 3.0 | 3 | 1.5 | 0.002 | 5,000 | 7,748 | 6,246 |
| Opisthokonta | Metazoa | Spiralia | 2 | 49 | 26.5 | 1 | 1.0 | - | 5,300 | 6,675 | 5,988 |
| Opisthokonta | Metazoa | Nemertea | 2 | 4 | 3.0 | 1 | 1.0 | - | 6,413 | 6,415 | 6,414 |
| Opisthokonta | Metazoa | Priapulida | 1 | 1 | 1.0 | 1 | 1.0 | - | 7,473 | 7,473 | 7,473 |
| Opisthokonta | Metazoa | Acanthocephala | 1 | 46 | 46.0 | 15 | 15.0 | 0.001 | 6,064 | 10,883 | 10,776 |
| Opisthokonta | Opisthokonta X | Opisthokonta XX | 1 | 1 | 1.0 | 1 | 1.0 | - | 5,946 | 5,946 | 5,946 |
| SAR | Alveolata | Apicomplexa | 151 | 59 | 3.3 | 30 | 1.6 | 0.003 | 4,647 | 10,794 | 6,338 |
| SAR | Alveolata | Ciliophora | 28 | 3 | 1.1 | 1 | 1.0 | - | 5,209 | 6,274 | 5,584 |
| SAR | Alveolata | Dinoflagellata | 7 | 33 | 11.0 | 13 | 4.3 | 0.010 | 5,078 | 5,844 | 5,509 |
| SAR | Alveolata | Perkinzoa | 2 | 2 | 1.5 | 1 | 1.0 | - | 5,936 | 6,044 | 5,990 |
| SAR | Rhizaria | Cercozoa | 3 | 6 | 5.3 | 2 | 1.3 | 0.002 | 5,107 | 6,545 | 6,118 |
| SAR | Rhizaria | Endomyxa | 1 | 1 | 1.0 | 1 | 1.0 | - | 4,702 | 4,702 | 4,702 |
| SAR | Stramenopiles | Oomycota | 157 | 72 | 9.0 | 15 | 1.8 | 0.008 | 4,261 | 7,317 | 6,054 |
| SAR | Stramenopiles | Ochrophyta | 28 | 376 | 17.6 | 7 | 1.5 | 0.003 | 4,799 | 6,383 | 5,897 |
| SAR | Stramenopiles | Bacillariophyta | 27 | 24 | 5.7 | 13 | 2.3 | 0.003 | 4,713 | 6,401 | 5,795 |
| SAR | Stramenopiles | Bigyra | 15 | 102 | 11.3 | 87 | 9.6 | 0.003 | 4,436 | 6,489 | 5,664 |
| SAR | Stramenopiles | Oochrophyta | 2 | 48 | 24.5 | 2 | 1.5 | 0.004 | 5,262 | 6,164 | 5,554 |

### Clustering analyses of sub-regions

To compare the level of molecular variation for the regions of the ribosomal operon often used in metabarcoding studies (Santoferrara, 2019), we extracted different parts (18S, 28S, V4, ITS1, and ITS2) from the entire operons and clustered these individually. To retrieve the regions, we first built an alignment of the operon variants using MAFFT. Next, we marked the boundaries of each region based on a set of primers and the identity score for the alignment. The primers used to identify the V4 region were 3NDf (Cavalier-Smith et al., 2009), and TAReukREV3 (Stoeck et al., 2010); the end of 18S (which is also the start of ITS1) was confirmed by EukB (Medlin et al., 1988), while the primer fITS7 was used to split ITS1 and ITS2 in 5.8S (Ihrmark et al., 2012). Finally, the beginning of 28S/end of ITS2 was confirmed with the primer ITS4 (White et al., 1990) (Fig. 1)

**Figure 1.**
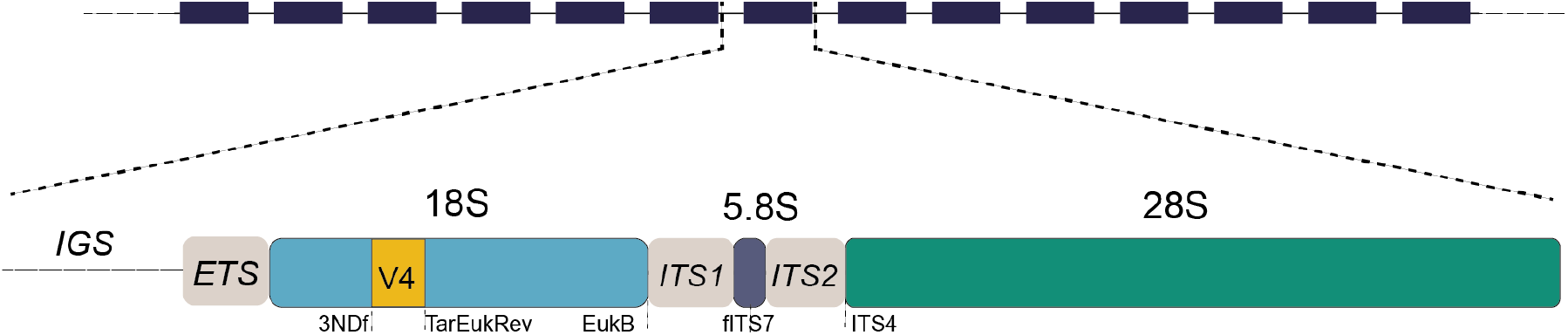
Typical arrangement of the ribosomal operon in eukaryotes. The ribosomal operon is commonly arranged in tandem repeats with intergenic spacers (IGS) between copies and an external transcribed spacer located before the 18S gene. The genes 18S, 5.8S, and 28S are separated by the internally transcribed spacers (ITS1 and ITS2). The binding sites of primers used to separate regions of interest for the clustering analyses are annotated underneath the operon.

In addition to the 18S, 28S, ITS1, ITS2, and V4 subregion, a superregion encompassing ITS1-5.8S-ITS2 was also extracted and clustered. Full intragenomic operon centroids and subsectioned datasets were processed separately. Following an initial cleanup to remove sequences containing more than 20 consecutive N-bases in any subregion, the sequences were dereplicated and clustered using VSEARCH (Rognes et al. 2016). Clustering was performed from 95.0% to 100% with 0.5 increments to see the effect of different clustering levels for each subregion (Bonin et al., 2023).

### Choosing a representative sequence

As a final resource, we chose one representative sequence from each genome assembly with the highest copy number of all the operon variants from a genome if more than one variant existed. In a few genome assemblies (68), two or more variants with the highest copy number were equally abundant, and they were not included in the reduced reference dataset in ROD.

### Data visualization

Visualization of the number of genomes, operon variants, copy numbers, and genetic distance on the eukaryotic Tree of Life was done with the package *metacoder* (Foster et al., 2017) in R v4.3.3 (R Core Team, 2024). The ridgeplots were made in R with *ggplot2* (Wickham, 2016) and *ggridges* (Wilke, 2024), and the colors were based on the package *wesanderson* (Ram & Wickham, 2023).

## Results

While a signal for the ribosomal operon was found in 17,107 out of the 34,701 genome assemblies, full-length rDNA operon(s) were detected in 11,935 genome assemblies (34.4%). The operons were clustered intragenomically at 99.0% sequence similarity to remove tentative technical errors from sequencing and assembly, as well as to record operon variants diverging by more than 1% in sequence identity. The number of clusters represents the number of operon variants per genome, and the size represents the number of copies found for a given variant. The set of operon variants (69,480), as well as a reference set including only the most abundant operon variant per genome (11,867), are available in the ROD database (https://github.com/krabberod/ROD). The release used in this study is ROD v1.0 (GoldenRod), which is also deposited on Zenodo (https://zenodo.org/records/10951604).

Figure 2 shows the distribution of the detected operons across major taxonomic groups. Fungi, which are intensively sequenced partly due to their small genomes, represent 55.5% of the genomes in ROD, followed by Metazoa (22.7%) and Viridiplantae (17.0%) (Fig. 2, Table 1). A single copy of the operon was found in 5,601 (46.9%) of the assemblies, while 53.1% had multiple copies. Among the genomes with multiple copies, most had between 2-10 copies (3,385), 2,241 had between 11 and 100 copies, 708 had more than 100 copies, and 92 genomes had more than 1,000 copies (Fig. 2). Most copies were observed in genome assemblies of Viridiplantae (Fig. 2, Table 1). The highest copy number, 5,947, was found in the genome of *Zea mays* (genome assembly id GCA_024505845 (Zhao et al., 2022)). Also, many Metazoa possessed several copies (Table 1), with the insect *Clusia tigrina* having the highest number with 4,691 copies (genome assembly id GCA_920105625 (Crowley et al., 2023)).

**Figure 2.**
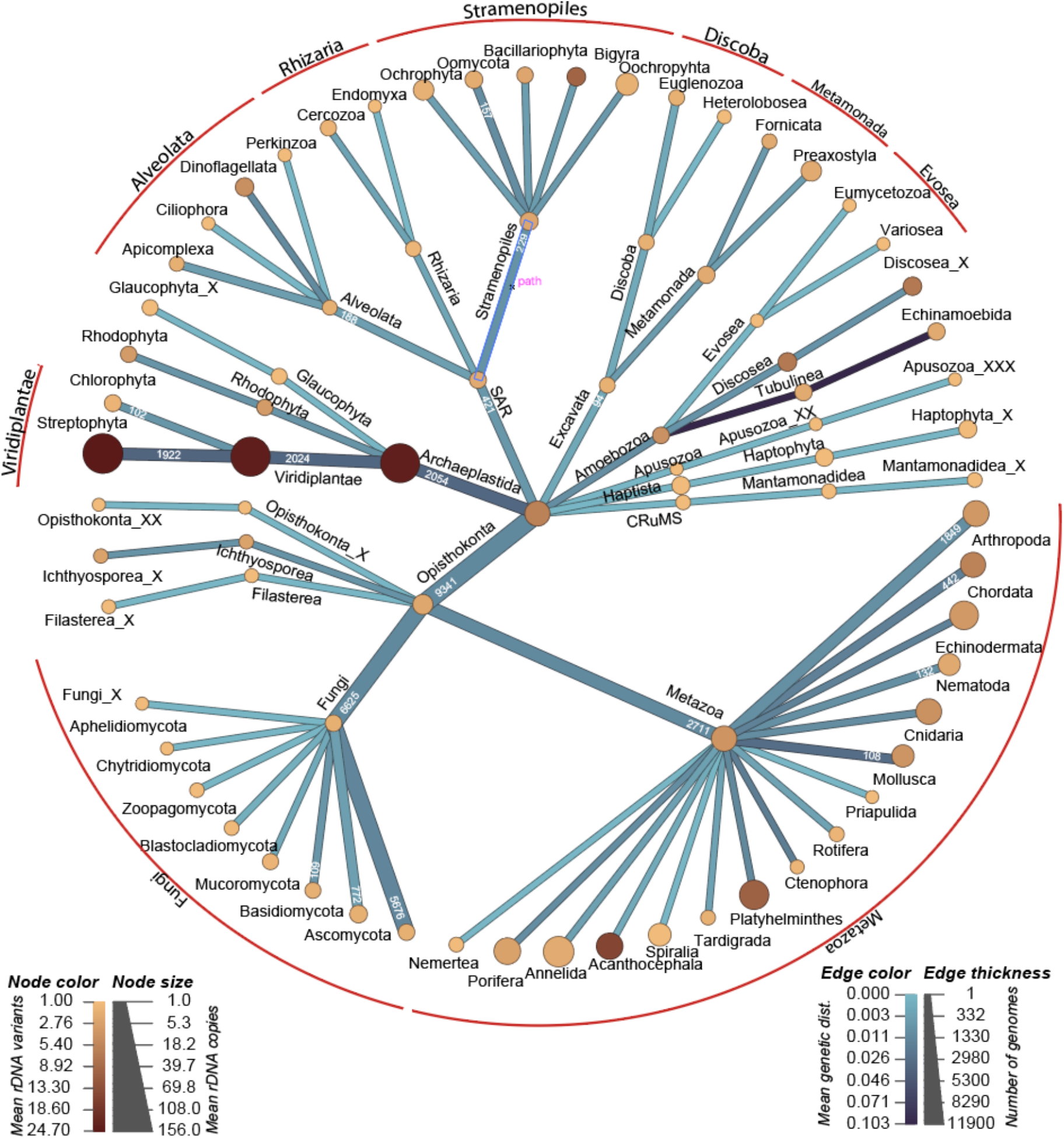
The Distribution of the 69,480 analyzed rDNA operons across the eukaryotic tree of life in the Ribosomal Operon Database (ROD). The unrooted tree is a schematic representation of eukaryotes produced using Metacoder (Foster et al., 2017). The thickness of the branches represents the number of genomes for the clade, and the color of the branches is the weighted genetic distance for the variants. The size of the nodes represents the Mean rDNA copy number for the genomes of that node, and the color of the nodes represents the mean number of variants.

The number of operon variants per genome varied considerably between major taxonomic groups (Fig. 2). While 8,356 out of the 11,935 genome assemblies with a full-length operon had a single operon variant, 2,733 had between 2-10 variants, 730 between 11 and 100 variants, and 117 had more than 100 variants. As with the copy numbers, Viridiplantae included the most variant numbers (Fig. 2). Two plant species were the only genomes with more than 1000 operon variants: *Thlaspi arvense* with 1,017 variants (genome assembly id GCA_018983045 (Geng et al., 2021) and *Silybum marianum* with 2,097 variants (genome assembly id GCA_001541825.1 (Shim et al., 2023) (Table 1 and Supplementary table 1). In contrast, fungal genome assemblies generally had fewer variants; the highest number was 81, found in the ascomycete *Leptosphaeria maculans* (genome assembly id GCA_022343315 (Tian et al., 2024)).

To chart the amount of intragenomic variation, we calculated the genetic distance between all operon copies for each assembly separately (Fig. 2). Among the most abundant taxonomic groups sampled, Viridiplantae possessed the highest average genetic distance between copies (0.033). In comparison, the different sub-groups in Metazoa and Fungi had, on average, between 0.001-0.027 (Metazoa) and 0-0.12 (Fungi) sequence distance between copies (Table 1). The highest genetic divergence for a single assembly was observed in the assembly of the fungi *Neurospora* sp. CHS-2018c. Here, we found 30 operon variants, all single copies, with an average genetic distance of 0.215 (supplementary table 1).

The length of the operon varied extensively across the organismal groups (Fig. 3). The shortest operon found was 4,136 bp in the plant *Capsicum annuum* (assembly id GCA_030864225 (Estrada et al., 2024)), and the longest was 16,463 bp in the large bee-fly *Bombylius major* (assembly id GCA_932526495 (Lawniczak et al., 2023)). However, 86.0% of all operons had a length between 5,000 and 6,500 bp, and the median length was 5,779 bp (Fig. 3). Most of the operons in Fungi and Viridiplantae ranged between 5,500-6,100 bp in length (Fig. 3). The length variation was more extensive in Metazoa, where some groups, like Acanthocephala with an average operon length of 10,776 bp, possessed long operons.

**Figure 3.**
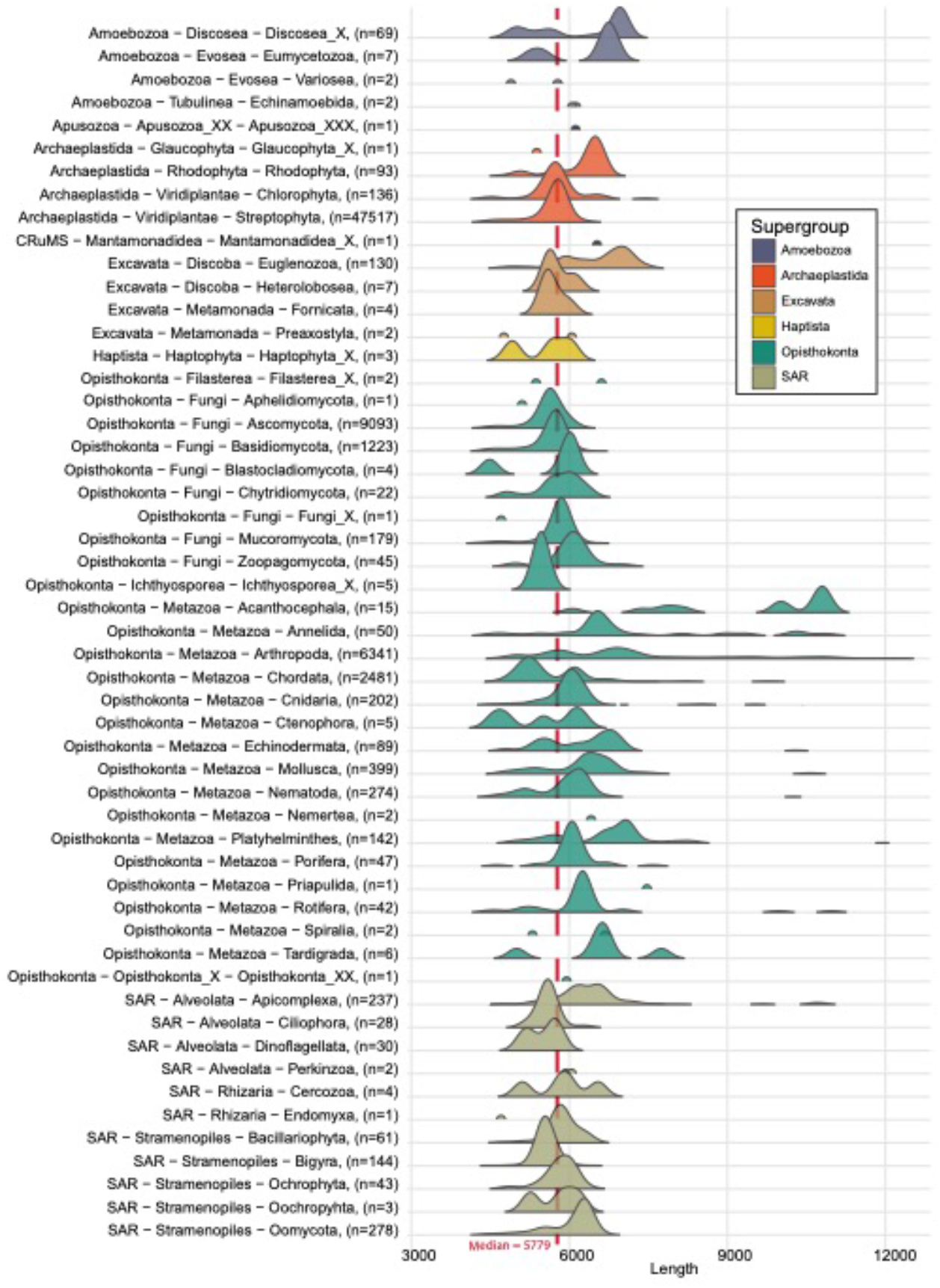
Length distribution of the recorded rDNA operons across the eukaryotic subdivisions in the Ribosomal Operon Database (ROD). The ridge plot represents the density distribution of lengths within each group. The number of genomes for each subdivision is written in parentheses. The median length is recorded as a red dashed line.

To investigate the level of molecular variation across full-length operons compared to the 18S and 28S regions, as well as the shorter V4, ITS, ITS1, and ITS2 regions, and to understand how this variation influences cluster numbers, these regions were separately clustered across various sequence identity thresholds (Fig. 4a). Across all clustering thresholds, the 18S and V4 regions yielded the fewest clusters. In contrast, the more variable ITS region and the full-length operons produced the highest numbers (Fig. 4a).

**Figure 4.**
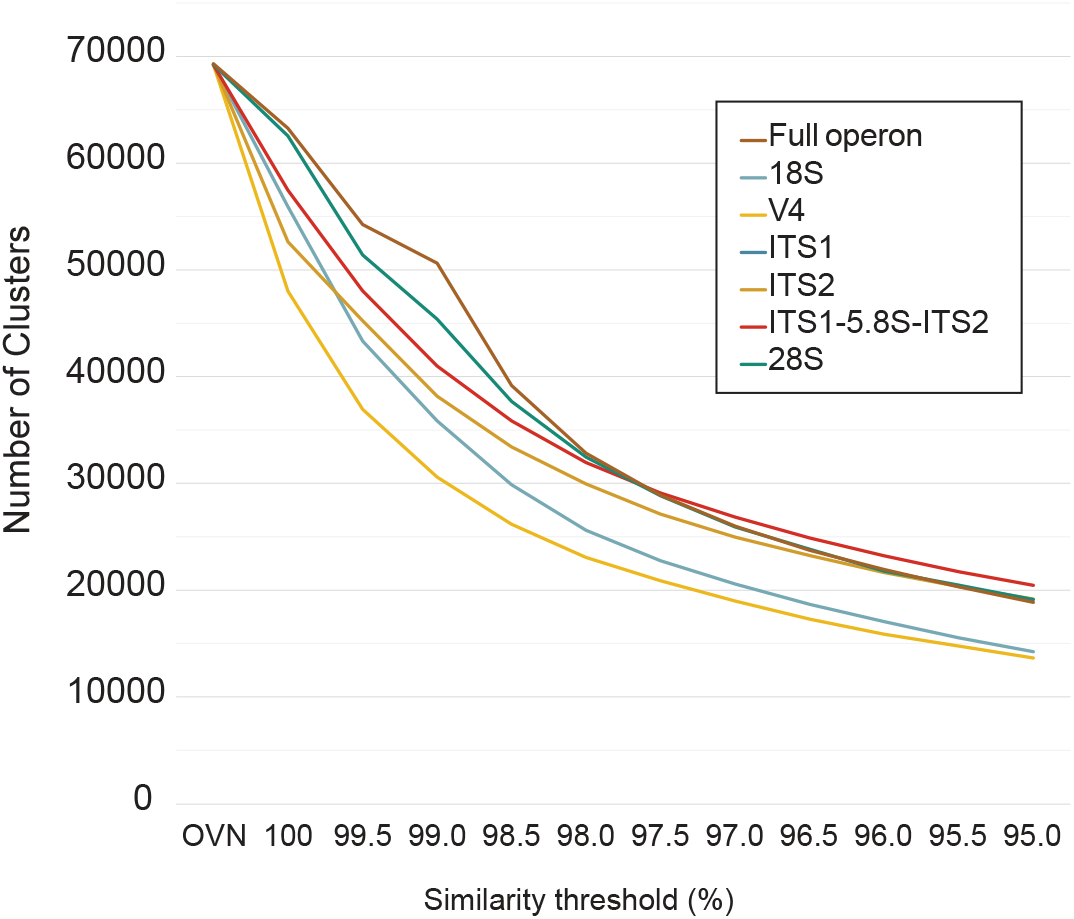
Cluster analysis of the 69,480 full-length operon variants in ROD and different sub-parts commonly used in DNA metabarcoding studies. The number of sequence clusters (y-axis) plotted against the clustering threshold for the entire operon, as well as the different subparts. On the x-axis, OVN is the original variant number, i.e., the number of operons before clustering (69,480), while 100% gives the number of variants after collapsing identical operon variants across all genomes, similar to a dereplication.

## Discussion

Recently, long-read sequencing approaches have been adopted in microbial ecology and DNA metabarcoding to obtain more detailed phylogenetic information about the microbial communities under investigation (Jamy et al., 2020, 2022; Tedersoo et al., 2020). More solid phylogenetic information is especially beneficial for poorly studied organisms that can be hard to place into a broader phylogenetic context (Krabberød et al., 2011). However, for long-read DNA metabarcoding analyses, we need reference sequences that cover longer stretches of the rDNA region, preferably the entire operon. Motivated by this, we provide full-length rDNA operon reference sequences assembled into the Ribosomal Operon Database (ROD). In addition to functioning as reference sequences for taxonomic annotation, the sequences in the ROD database may also fulfill other needs, such as reference sequences during chimera-checking, primer design, or phylogenetic binning. ROD currently includes 69,480 operon variants and their copy numbers extracted from 11,935 genomes. In addition, a reference set with only one operon (i.e., the most frequent) per genome is made available, which might be more useful in some investigations.

The reference sequence data have undergone multiple error correction steps, where contaminant and tentatively spurious sequences have been parsed out. Nevertheless, we cannot dismiss the possibility of spurious operons in the database. For example, we observed that many of the genome assemblies (29.3% of the full-length operons) were contaminated by other organisms. However, in this study, we only report on the observed operon variability in existing genome assemblies, and we do not aim to separate the different sources of variability.

We argue that full-length rDNA operon reference sequences should not be established based on concatenated sequences from different sub-parts, e.g., 18S and 28S, derived from multiple specimens, as this could lead to erroneous concatenation of different operon variants. Moreover, high-quality reference sequences should preferably be obtained from identified reference material, such as the sequencing of the full-length operon or genome from individuals in the case of multicellular organisms or axenic cultures in the case of single-celled organisms. Reference sequences obtained from environmental sequence data that have been assigned a taxonomy based on other reference sequences can also be useful but should be treated with caution since this represents indirect evidence. Moreover, a ‘nearest neighbor’ approach, in which the query sequence is assumed to have the same taxonomy as the closest match in a database, is highly dependent on the presence of a close match in the database or the training set (Murali et al., 2018). Adding to the dilemma, it is not always given how closely a sequence must match the reference in order to be correctly assigned to a given taxonomic rank, and the temptation to use the same naïve similarity cut-off for different groups of eukaryotes can easily lead to wrongful taxonomic annotations. In such cases, errors can quickly spread and multiply.

The copy number of the ribosomal operon can vary substantially between lineages of eukaryotes with as many as 19,300 reported in animals, more than 26,000 in plants (Prokopowich et al., 2003) and up to 1,442 in fungi (Lofgren et al., 2019). There are much fewer genomes available from microbial eukaryotes (protists). It has been shown, however, that many protist groups have a huge number of copies. In Foraminfera, the average number varies between 4,000 and 50,000 in different species (Milivojević et al., 2021), while the ciliate *Halteria grandinella* has as many as 567,893 copies (Wang et al., 2017). The number of copies can even vary between individuals of the same species, and this variation in copy number has been reported to result from both genetic and environmental factors (Lavrinienko et al., 2021; Milivojević et al., 2021; Symonová, 2019). Reports on copy number variation are mostly done with qPCR or other cytological measurements, and such high numbers seldom occur in genome assembly projects. The reason is that until recently, repeated elements were very hard to resolve accurately due to the short read length of the sequencing platforms (Tørresen et al., 2019). In genome assemblies based on short reads, repeated elements tend to collapse into one representative sequence, and the number of actual copies per genome reported in the current version of ROD is, therefore, likely an underestimate. With the advent of highly accurate long-read sequencing technologies, the possibility of resolving long repeats is increasing and will most likely improve in the future.

It is commonly known that copies are usually found in tandem repeats, and concerted evolution mechanisms keep the copies highly uniform (Eickbush & Eickbush, 2007). However, multiple operon variants, here defined as operons diverging by more than 1% in sequence identity, were discovered in approximately 30% of the genomes analyzed. Especially in the analyzed plant genomes, we observed a high number of operon variants per genome, which could be due to their often polyploid nature. But even in haploid genomes, multiple variants can be expected if the multiple operon copies are not homogenized by concerted evolution (Bradshaw et al., 2023). The occurrence of multiple, divergent operon variants per genome (or individuals) can lead to inflation of the diversity (Lavrinienko et al., 2021; Thornhill et al., 2007). Operons can occasionally appear in genome assemblies as multiple variants for technical reasons. Our analysis showed that more than 50% of the genomes from NCBI had ribosomal operons from contaminating species. These can readily be removed by blasting and comparing the taxonomic affiliation of the sequence to the expected taxonomy of the genome being assembled. These issues can be difficult to resolve, but as sequencing platforms improve and sequences become longer and more accurate, these problems are expected to be improved. Thanks to advancements in sequencing and bioinformatics methods, we envision that future genome assemblies will be of even higher quality.

Using the full ROD dataset as reference sequences for environmental sequencing studies can help address the issue of multiple operon variants during the taxonomic annotation process. This might be especially important for organismal groups with divergent operons, such as Viridiplantae. Due to the initial 99% clustering of operons within each genome, we may underestimate the true variant numbers.

High variation in the operon copy number was observed across the eukaryotic tree of life, with an average high copy number present in Viridiplantae. The copy number variability will introduce PCR bias in DNA metabarcoding projects with environmental samples, where genomes and taxa with high copy number will be preferentially amplified (Lavrinienko et al., 2021). If the copy number variability becomes better known, measures may be taken to account for this bias in the final data (Stoddard et al., 2015)

Significant operon length differences are present between and within various taxonomic groups. Based on our data, the length variability seems especially prominent in Metazoa. One major cause for the extremely long variants is the presence of introns in several lineages (Bhattacharya et al., 2000; Chang et al., 2017; Hedberg & Johansen, 2013; Karpov et al., 2017). During the amplification of longer stretches or the entirety of the rDNA operon, the length differences will introduce PCR bias, where taxa with shorter variants will be preferentially amplified. In theory, if the length variation is known across organismal groups, the bias could be modeled and taken into account.

Due to the varying rates of molecular evolution across the rDNA operon, different regions provide different phylogenetic and taxonomic resolutions. This variation is further complicated by differing evolutionary rates in rDNA operons across the Tree of Life, making the selection of optimal barcoding regions and clustering settings for operational taxonomic units (OTUs) a significant challenge (Phillips et al., 2022). For example, while the ITS region frequently offers species-level resolution for fungi (Schoch et al., 2012), using more conserved markers such as V4 often groups related species into the same OTUs at the same clustering level. Our clustering analyses show that both the ITS regions and the entire operon provide more fine-grained taxonomic resolution than 18S and the hypervariable region V4. These findings indicate the necessity for marker-specific bioinformatic processing strategies, including adjusting clustering levels to avoid potential over-splitting or merging sequences during OTU construction (Bonin et al., 2023; He et al., 2015).

The ROD database currently possesses a taxonomically highly skewed collection of reference sequences that reflect the distribution of available genomes in NCBI. Fungi are the dominating group, followed by Metazoa and Viridiplantae. Due to their small genome size and biotechnological importance, many fungal genomes have been sequenced and are available in public databases (de Vries et al., 2018; Xu, 2023). In the future, we expect more genomes from neglected branches on the Tree of Life distribution, including more protist genomes. Due to advances in single-cell genomics, genomes from single-celled protists will expectedly become more available as references (Sieracki et al., 2019). Because of the steadily increasing pace of genome sequencing, we expect a strong increase in the number of operons that will become available for inclusion in ROD in the near future. We intend to provide regularly updated versions of ROD.

## Acknowledgments

We would like to thank Ane-Rikke N. Oenting for invaluable feedback, the University of Oslo for financial support, and the National Infrastructure for High-Performance Computing in Norway (Sigma2) for access to computational resources on projects NN9338K, NN9725K, and NN9699K.

## Data Accessibility and Benefit-Sharing

The full set (69,480) and the reference set of operon variants (11,867) are available in the ROD database (https://github.com/krabberod/ROD). The release used in this study is ROD v1.1 (GoldenRod), which is also deposited on Zenodo (https://zenodo.org/records/11060492).

## Author Contributions

AKK conceived the idea of establishing ROD and assembled the database. AKK and EES did the analyses, HK and AKK drafted the manuscript while all co-authors commented and edited.

